# Molecular basis for the maintenance of lipid asymmetry in the outer membrane of *Escherichia coli*

**DOI:** 10.1101/192500

**Authors:** Jiang Yeow, Kang Wei Tan, Daniel A. Holdbrook, Zhi-Soon Chong, Jan K. Marzinek, Peter J. Bond, Shu-Sin Chng

## Abstract

A distinctive feature of the Gram-negative bacterial cell envelope is the asymmetric outer membrane (OM), where lipopolysaccharides (LPS) and phospholipids (PLs) reside in the outer and inner leaflets, respectively. This unique lipid asymmetry renders the OM impermeable to external insults. In *Escherichia coli*, the OmpC-MlaA complex is believed to maintain lipid asymmetry by removing mislocalized PLs from the outer leaflet of the OM. How it performs this function is unknown. Here, we define the molecular architecture of the OmpC-MlaA complex to gain insights into its role in PL transport. We establish that MlaA sits entirely within the bilayer in complex with OmpC and provides a hydrophilic channel possibly for PL translocation across the OM. Furthermore, we show that flexibility in a hairpin loop adjacent to the channel modulates MlaA activity. Finally, we demonstrate that OmpC plays an active role in maintaining OM lipid asymmetry together with MlaA. Our work offers glimpses into how the OmpC-MlaA complex transports PLs across the OM and has important implications for future antimicrobial drug development.

## Introduction

The outer membrane (OM) of Gram-negative bacteria is an extremely asymmetric bilayer, comprising lipopolysaccharides (LPS) in the outer leaflet and phospholipids (PLs) in the inner leaflet (1, 2). LPS molecules pack tightly together in the presence of divalent cations to form an outer layer with markedly reduced fluidity and permeability (3). Thus, the OM serves as an effective barrier against toxic compounds including detergents and antibiotics. This function is fully dependent on the establishment and maintenance of lipid asymmetry; cells generally become more sensitive to external insults when OM lipid asymmetry is disrupted, which is typically characterized by the accumulation of PLs in the outer leaflet (4, 5). The OM is also essential for viability.

The requisite lipid asymmetry of the OM is likely initially established by direct placement of LPS and PLs into the outer and inner leaflets, respectively. LPS assembly into the outer leaflet of the OM is mediated by the well-established Lpt (lipopolysaccharide transport) machinery (6), but proteins that transport and insert PLs into the inner leaflet have not been identified. For entropic reasons, there is a natural tendency for PLs to appear in the outer leaflet of the OM, although how they traverse the bilayer is unclear. This occurs more readily with perturbations in the OM, especially when assembly of other OM components is disrupted (4, 5, 7). Since loss of lipid asymmetry compromises the barrier function of the OM, several mechanisms exist to remove PLs aberrantly localized in the outer leaflet of the membrane: (i) the OM phospholipase OmpLA hydrolyzes both acyl chains from outer leaflet PLs (8), (ii) the OM acyltransferase PagP transfers an acyl chain from outer leaflet PLs to LPS (9) or phosphatidylglycerol (PG) (10), and (iii) the OmpC-Mla system, a putative PL trafficking pathway, removes outer leaflet PLs and shuttles them back to the inner membrane (IM) (11, 12).

The OmpC-Mla system comprises seven proteins located across the cell envelope. Removing any component results in PL accumulation in the outer leaflet of the OM, and therefore sensitivity to SDS/EDTA (11, 12). The OM lipoprotein MlaA forms a complex with osmoporin OmpC that likely extracts PLs from the outer leaflet of the OM (12). The periplasmic protein MlaC serves as a lipid chaperone and is proposed to transport lipids from the OmpC-MlaA complex to the IM (11, 13, 14). At the IM, MlaF and MlaE constitute an ATP-binding cassette (ABC) family transporter together with two auxiliary proteins, MlaD and MlaB (14, 15); this complex presumably receives PLs from MlaC and inserts them into the membrane. MlaD has been shown to bind PLs, while MlaB is important for both assembly and activity of the transporter (15). Recently, the function of the OmpC-Mla system in retrograde (OM-to-IM) PL transport has been demonstrated in *E. coli* (7).

The molecular mechanism by which the OmpC-MlaA complex extracts PLs from the outer leaflet of the OM, presumably in an energy-independent manner, is an interesting problem. Aside from the LptDE machine, which assembles LPS on the surface (4, 5), the OmpC-MlaA complex is the only other system proposed to catalyze the translocation of lipids across the OM. OmpC is a classical trimeric porin that typically only allows passage of hydrophilic solutes across the OM (3, 16), while MlaA is believed to be anchored to the inner leaflet of the membrane; how the two proteins are organized in a complex for the translocation of amphipathic PLs is not known. In this paper, we establish that MlaA is in fact an integral membrane protein that forms a channel adjacent to OmpC trimers in the OM, likely allowing the passage of PLs. We first demonstrated that MlaA binds the OmpC trimer within the OM bilayer by mapping the interaction surfaces using in vivo crosslinking. Using a recently predicted structural model of MlaA (17), we obtained molecular views of the OmpC-MlaA complex by molecular dynamics (MD) simulations, and experimentally established the existence of a hydrophilic channel within OM-embedded MlaA. Combining charge mutations in this channel modulated MlaA activity, suggesting functional importance. Furthermore, mutations altering the flexibility of a hairpin loop that could interact with the hydrophilic channel led to predictable in vivo effects on MlaA function. Finally, we identified a key residue on OmpC found at the OmpC-MlaA interacting surface that is important for proper function of the complex. Our findings provide important mechanistic insights into how PLs may be translocated across the OM to ensure proper lipid asymmetry.

## Results

### The OmpC trimer contacts MlaA directly along its membrane-facing dimeric interfaces

To develop a detailed architectural understanding of the OmpC-MlaA complex, we carried out in vivo photocrosslinking to map intermolecular interactions within the complex. Guided by the crystal structure of the OmpC trimer (18), we introduced the UV-crosslinking amino acid, *para*-benzoyl-L-phenylalanine (*p*Bpa), at 49 positions in OmpC via amber suppression (19). Initial selection focused on residues that are either solvent-accessible (i.e. loop and lumen) or located near the membrane-water boundaries (i.e. aromatic girdle). Upon UV irradiation, a ˜65 kDa crosslinked band that contains both OmpC (˜37 kDa) and MlaA (˜28 kDa) could be detected in cells expressing OmpC variants substituted with *p*Bpa at three positions (L50, Q83, or F267) (Fig. 1*A*). These residues are found on the periplasmic turns at the dimeric interfaces of the OmpC trimer (Fig. 1*C*), thus localizing possible binding sites for MlaA. We have previously proposed that OmpC may allow MlaA to traverse the bilayer and gain access to PLs that have accumulated in the outer leaflet of the OM (12). As none of the six selected residues in the OmpC lumen crosslinked to MlaA, we decided to probe for interactions between MlaA and the membrane-facing side walls of OmpC, specifically around the dimeric interfaces of the OmpC trimer. Remarkably, out of the additional 49 positions tested in this region, 10 residues allowed photoactivated crosslinks between OmpC and MlaA when replaced with *p*Bpa (Fig. 1*B* and Fig. S1). In total, these 13 crosslinking residues clearly demarcate an extensive MlaA-interacting surface on OmpC (Fig. 1*C*). This explains why OmpC exhibits strong interactions and can be co-purified with MlaA on an affinity column, as we have previously reported (12). Two positions, Y149 and L340, are located right at or near the membrane-water boundary exposed to the extracellular environment, suggesting that MlaA traverses the entire width of the OM. We conclude that MlaA binds along the dimeric interfaces of the OmpC trimer in the membrane.

**Figure 1.**
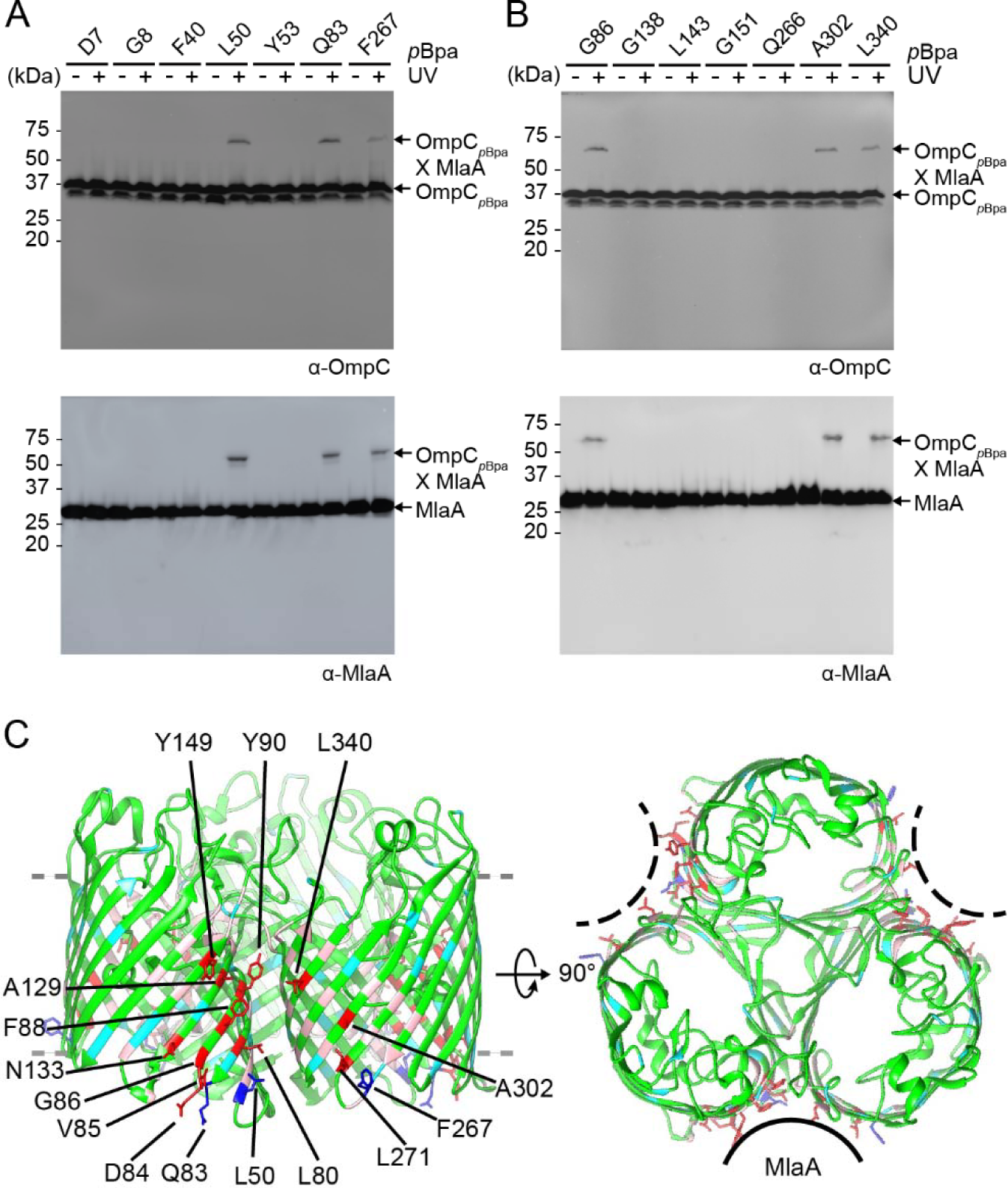
MlaA binds at the dimeric interfaces of the OmpC trimer in vivo. (*A*, *B*) Representative immunoblots showing UV-dependent formation of crosslinks between OmpC and MlaA in Δ*ompC* cells expressing OmpC substituted with *p*Bpa at indicated positions, selected in a (*A*) global, or (*B*) localized search. (*C*) Side (*left*) and top (*right*) views of cartoon representations of the crystal structure of *E. coli* OmpC (PDB ID: 2J1N) (18) with positions that crosslink to MlaA highlighted. Residues selected in the global search for MlaA interaction are colored *cyan* (no crosslinks) and *blue* (sticks; crosslinks detected), while those selected in the localized search are colored *light pink* (no crosslinks) and *red* (sticks; crosslinks detected). The OM boundary is indicated as gray dashed lines. MlaA binding sites are indicated as solid or dashed curves on the top-view representation.

### Two specific regions on MlaA contact the membrane-facing dimeric interfaces of the OmpC trimer

We sought to map in greater detail the OmpC-MlaA interacting surface in vitro. To do that, we first overexpressed and purified the OmpC-MlaA complex to homogeneity. We showed that this complex forms a single peak on size exclusion chromatography (SEC) (Fig. 2*A*). OmpC within this complex exhibits the characteristic heat-modifiable gel shift commonly observed for OM β-barrel proteins (20), consistent with the presence of the folded trimer. Multi-angle light scattering (MALS) analysis revealed that one copy of MlaA interacts with the OmpC trimer (Fig. S2), suggesting that only one of the three dimeric interfaces within the trimer is available for binding (Fig. 1*C*). We next performed protease digestion experiments to identify specific region(s) on MlaA that may interact stably with OmpC. OM β-barrel proteins such as OmpC are known to be protease-resistant (21). Given that some parts of MlaA contact OmpC within the membrane, we expect these bound regions to also be protected from proteolytic degradation. Treatment of the purified OmpC-MlaA complex with trypsin results in almost complete degradation of MlaA, with the OmpC trimer remaining intact (Fig 2*A*). Following SEC, however, we found that an ˜8 kDa peptide (presumably from MlaA) remains stably bound to the trimer. N-terminal sequencing and tandem mass spectrometry (MS) analyses revealed that this peptide corresponds to MlaA_D61-K124_ (Figs. 2*A* and 2*C*, and Fig. S3). These results suggest that MlaA interacts strongly with OmpC in the membrane in part via this specific region.

**Figure 2.**
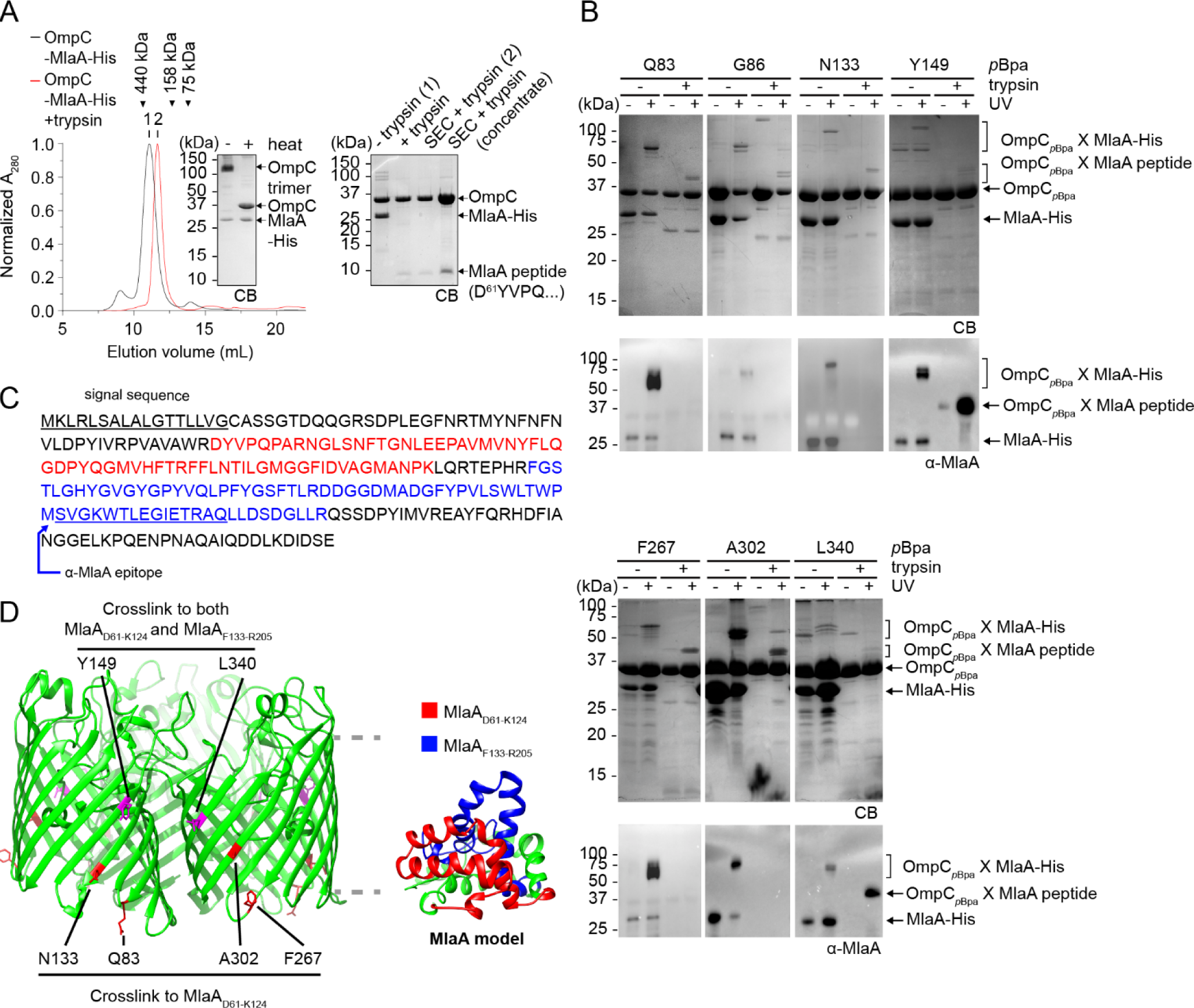
OmpC contacts two specific regions on MlaA. (*A*) SEC profiles and SDS-PAGE analyses of purified OmpC-MlaA-His complex before (*black*) or after (*red*) treatment with trypsin. Peak fractions from SEC were subjected to denaturing SDS-PAGE (15% Tris.HCl gel), followed by Coomassie Blue (CB) staining (*right*). Non-trypsin treated samples were also analysed by seminative SDS-PAGE (*left*). Edman degradation and tandem MS analyses revealed that the MlaA peptide that remains bound to OmpC following trypsin treatment begins at D61 (Fig. S3). (*B*) SDS-PAGE (15% Tris.HCl gel) and immunoblot analyses of purified OmpC*_p_*_Bpa-_MlaA-His complexes following sequential UV irradiation and trypsin digestion. The resulting OmpC*_p_*_Bpa_-MlaApeptide crosslinked products were N-terminally sequenced (see Fig. S3). (*C*) Amino acid sequence of MlaA with the two peptides found crosslinked to OmpC*_p_*_Bpa_ highlighted (*red*: MlaAD61-K124, *blue*: MlaAF133-R205). The signal sequence and α-MlaA binding epitope are underlined and annotated. (*D*) Cartoon representations of the crystal structure of *E. coli* OmpC with positions that crosslink to specific MlaA peptides indicated (*left*), and a structural model of MlaA (17) with peptides crosslinked by OmpC*_p_*_Bpa_ highlighted (*right*). The OM boundary is indicated as gray dashed lines.

To define how OmpC contacts the MlaA_D61-K124_ peptide, we next attempted to overexpress and purify *p*Bpa-containing OmpC variants in complex with MlaA, and determine which of the previously identified 13 OmpC residues interacts with MlaA_D61-K124_ in vitro. We sequentially performed UV crosslinking and trypsin digestion to potentially link MlaA_D61-K124_ to specific residues on OmpC. This approach may also allow trapping of other potential interacting regions of MlaA, which might not have been stably retained on the wild-type complex after trypsin digestion. We successfully detected trypsin-resistant crosslinked products for seven OmpC*_p_*_Bpa_-MlaA complexes (Fig. 2*B*); these appeared slightly above OmpC between 37 to 50 kDa, indicating that peptides in the range of ˜6-10 kDa are crosslinked to OmpC. N-terminal sequencing of six of the seven adducts showed the presence of OmpC, and an MlaA peptide beginning at residue D61 (Figs. 2*C* and 2*D*, and Fig. S3*C*). Given the approximate sizes of the crosslinked adducts, we concluded that all these residues interact with MlaA_D61-K124_. Interestingly, an additional peptide on MlaA starting at residue F133 was also found to crosslink at two (Y149 and L340) of these six positions on OmpC (Figs. 2*B*–*D*, and Fig. S3*C*). These adducts can be detected by an α-MlaA antibody that recognizes an epitope within V182-Q195 on MlaA (Fig. 2*B*), suggesting that a peptide from F133 to at least R205 (next trypsin cleavage site) may be crosslinked (Fig. 2*C*). Thus, in addition to MlaA_D61-K124_, our crosslinking strategy revealed a second point of contact (MlaAF133-R205) between OmpC and MlaA.

At the point of these findings, there was no available molecular structure for MlaA; however, a structural model has been predicted based on residue-residue contacts inferred from co-evolution analysis of metagenomic sequence data (17). Using a rigorously-validated quality score, this method of structure determination has generated reliable models for 614 protein families with currently unknown structures. We experimentally validated the model for MlaA by replacing residue pairs far apart on the primary sequence with cysteines, and showed that only those that are highly co-evolved (and predicted to be residue-residue contacts) allow disulfide bond formation in cells (Fig. S4). We therefore proceeded to use this MlaA model to understand the organization of the OmpC-MlaA complex. Interestingly, the positions of the two OmpC-contacting peptides on the MlaA model are spatially separated in a way consistent with the arrangement of the residues on OmpC that crosslink to each peptide (Fig. 2*D*). This not only reveals how MlaA may potentially be oriented and organized around the dimeric interface of the OmpC trimer, but also suggests that the entire MlaA molecule may reside in the membrane. In fact, the overall surface of MlaA, other than the putative periplasmic-facing region, is largely hydrophobic (Fig. S5). Furthermore, using all-atomistic MD simulations, we found that the structural fold of MlaA appears to be more stable in a lipid bilayer than in an aqueous environment (Fig. S6). Consistent with this, we note that even without its N-terminal lipid anchor, MlaA is not a soluble protein, and can only be extracted and purified from the OM in the presence of detergent. Collectively, these observations lend strong support to the validity of the predicted MlaA structure.

### MlaA provides a hydrophilic channel that may allow PL translocation across the OM

To obtain a physical picture of how OmpC interacts with MlaA within the complex, we used MD simulations to dock MlaA onto the OmpC trimer within a PL bilayer. Using a previously reported protocol (22), we first docked the MlaA model onto the OmpC trimer structure, both as rigid bodies. Interestingly, all initial docked structures contained MlaA binding at one dimeric interface of OmpC. Based on information derived from crosslinking, we selected two most consistent models differing slightly in how MlaA is oriented with respect to OmpC for unrestrained refinements using all-atomistic simulations. Multiple simulations were run for each MlaA orientation in a PL bilayer until overall root-mean-square deviations (RMSD) stabilized; remarkably, the resulting equilibrium models fulfilled all observed experimental crosslinking data. We performed clustering on all available trajectories, and identified the most populated conformations of the OmpC-MlaA complex in our simulations (Fig. 3*A* and Fig. S7*A* for one MlaA orientation, and Fig. 3*B* and Fig. S7*B* for the other). These conformational models all show MlaA sitting in the bilayer, tucked nicely into the dimeric interface of the OmpC trimer. Evidently, the MlaA_D61-K124_ peptide interacts extensively with OmpC in these models, consistent with why this peptide remains stably bound to OmpC after protease digestion (Fig. 2*A*). We also mutated several MlaA residues found at the OmpC-MlaA interfaces to *p*Bpa, and performed in vivo crosslinking experiments. We identified one position L109 that allowed strong photoactivatable crosslinking to OmpC when replaced with *p*Bpa (Fig. 3*C*). This residue lies within the MlaA_D61-K124_ peptide (Figs. 3*A* and 3*B*), confirming that this region does in fact contact OmpC in cells.

**Figure 3.**
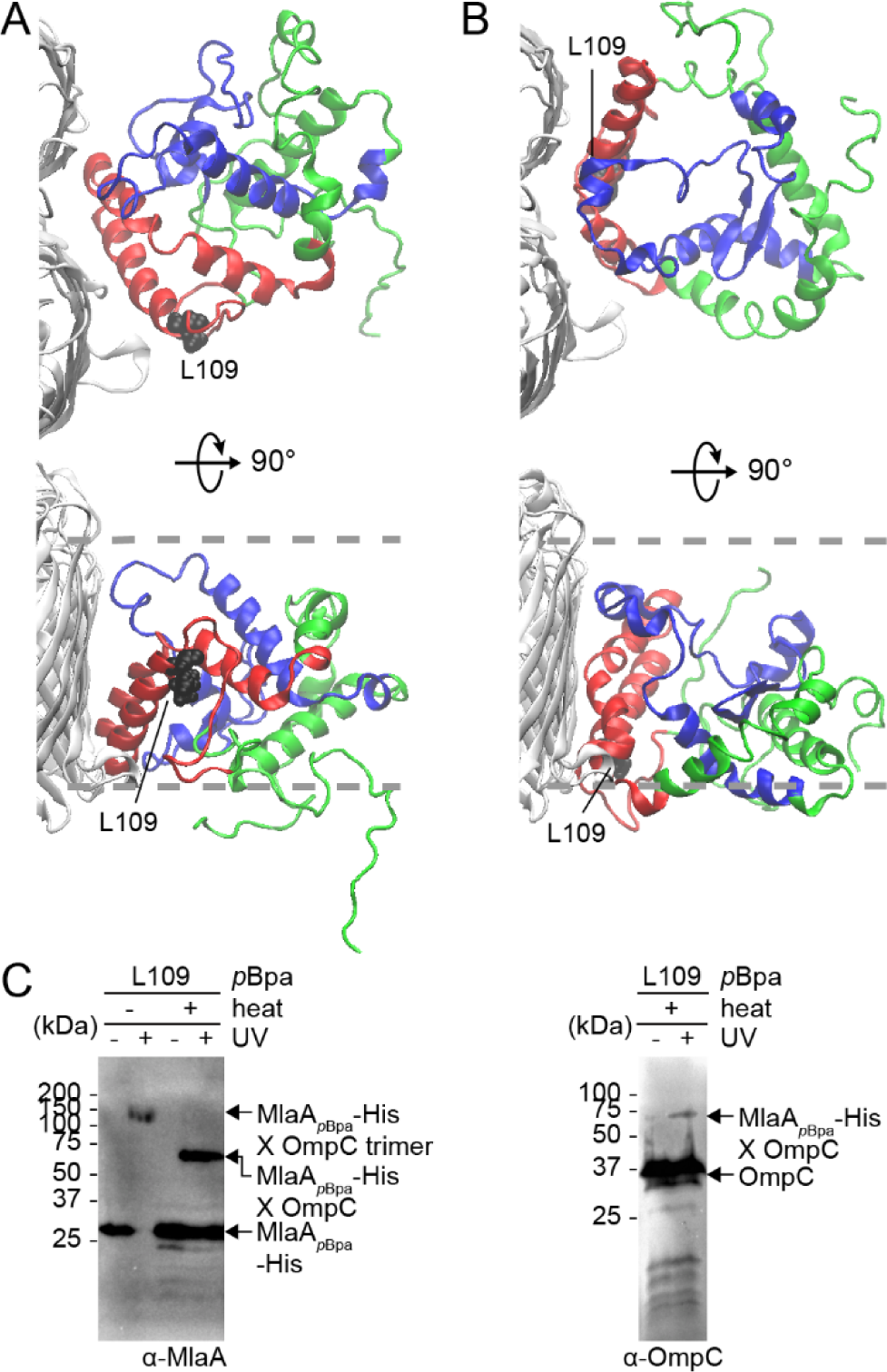
Molecular models of the OmpC-MlaA complex depict how MlaA may interact with OmpC in the OM bilayer. (*A*) and (*B*) Representative MlaA structures bound to OmpC in two possible orientations selected from all-atomistic MD simulation trajectories. MlaA_D61-K124_ and MlaAF133-R205 peptides are highlighted in *red* and *blue*, respectively, as in Fig. 2*D*. L109 is labelled and depicted as *black* spheres on MlaA. The OM boundaries are indicated as *gray* dashed lines. (*C*) Immunoblots showing UV-dependent formation of a crosslink between MlaA and OmpC in Δ*mlaA* cells expressing MlaA_L109_*_p_*_Bpa_-His. As expected, the crosslinked product also exhibits heat-modifiable gel shift, indicative of the presence of OmpC.

One striking feature present in all the simulated OmpC-MlaA structures is a negatively-charged hydrophilic channel within MlaA that spans the lipid bilayer (Fig. 4*A*, and Fig. S8). Based on its function in removing PLs from the outer leaflet of the OM, we hypothesize that this channel may allow passage of charged headgroups as PLs translocate across the membrane. To test the existence of this hydrophilic channel in MlaA in cells, we selected 27 residues in and around the putative channel in a representative model (Fig. 4, and Fig. S9), and determined their solvent accessibility using the substituted cysteine accessibility method (SCAM). We first showed that these cysteine mutants are functional (Fig. S9*C*). Solvent-exposed residues are expected to be reactive with the charged membrane-impermeable thiol-labelling reagent, sodium (2-sulfonatoethyl)methanethiosulfonate (MTSES). Remarkably, we found that residues predicted to be within the putative channel (Figs. 4*A* and 4*B*) or at the membrane-water boundaries (Figs. S9*A* and S9*B*) are indeed accessible to MTSES. We therefore conclude that MlaA forms a hydrophilic channel in the OM in cells.

**Figure 4.**
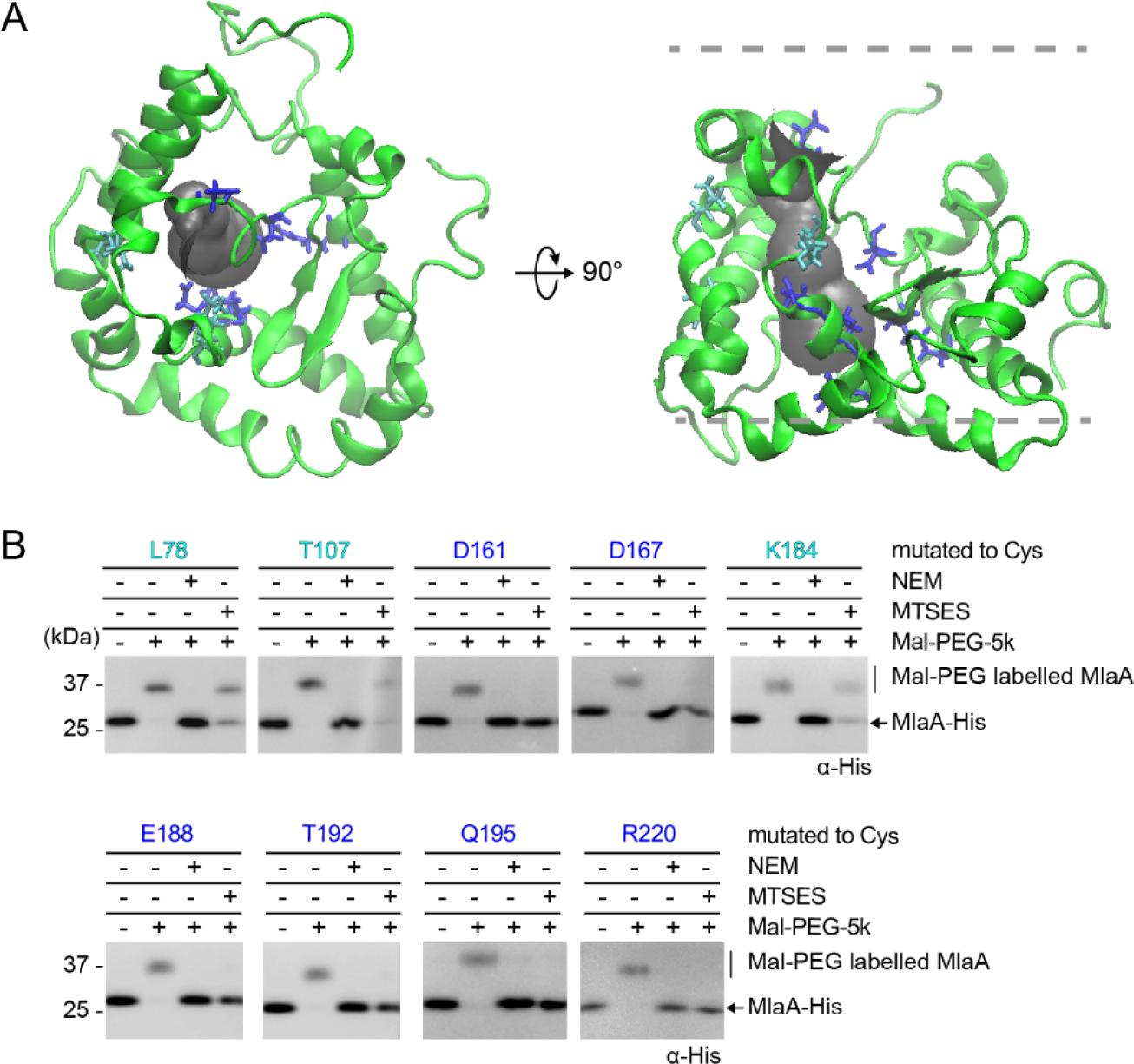
MlaA forms a hydrophilic channel in the OM. (*A*) A representative structure of MlaA from all-atomistic MD simulations with its putative channel depicted in *gray*. Residues in the channel that are fully or partially solvent accessible, based on SCAM in (*B*), are highlighted in *blue* and *cyan*, respectively. (*B*) Immunoblots showing maleimide-polyethylene glycol (MalPEG) alkylation of MlaA variants containing channel-facing residues substituted with cysteine (as depicted in (*A*)) following labelling by membrane permeable *N*-ethylmaleimide (NEM) or impermeable (MTSES) reagents. Mal-PEG alkylated MlaA_Cys_-His variants show a ˜5 kDa mass shift. Positions fully or partially blocked by MTSES, which reflects the level of solvent accessibility, are highlighted in *blue* or *cyan*, respectively.

To ascertain whether this channel is functionally important, we separately mutated 19 polar and (negatively) charged residues near or within the channel to alanine or arginine, and tested for MlaA function. However, all of these MlaA mutants are functional (Fig. 5*B*, and Fig. S10*A*), indicating that single residue changes are not sufficient to perturb channel properties to affect PL transport. We noticed that four (D160, D161, D164, and D167) of the five negatively-charged channel residues are in close proximity (Fig. 5*A*). To alter channel properties more drastically, we combined arginine mutations for these aspartates; interestingly, only the D161R/D167R double mutant (Fig. S10*B*) and D160R/D161R/D164R triple mutant (MlaA_3D3R_) disrupted function; cells expressing these variants are highly sensitive to SDS/EDTA (Fig. 5*B*, and Fig. S10*B*). For the MlaA_3D3R_ mutant, this is likely due to the accumulation of PLs in the outer leaflet of the OM (as judged by PagP-mediated acylation of LPS; Fig. 5*C*). In fact, this mutant exhibits OM defects that are more pronounced than the Δ*mlaA* strain (Fig. 5*B*), suggesting gain of function. Consistent with this idea, the *3D3R* mutation gives rise to the same defects in strains also expressing the wild-type *mlaA* allele, revealing a dominant negative phenotype (Figs. 5*B* and 5*C*). We showed that MlaA_3D3R_ is produced at levels comparable to wild-type MlaA on a plasmid (Fig. S11*A*), and is still able to interact strongly with OmpC (Fig. S11*B*). Taken together, these results suggest that the MlaA channel plays a functional role in maintenance of lipid asymmetry.

**Figure 5.**
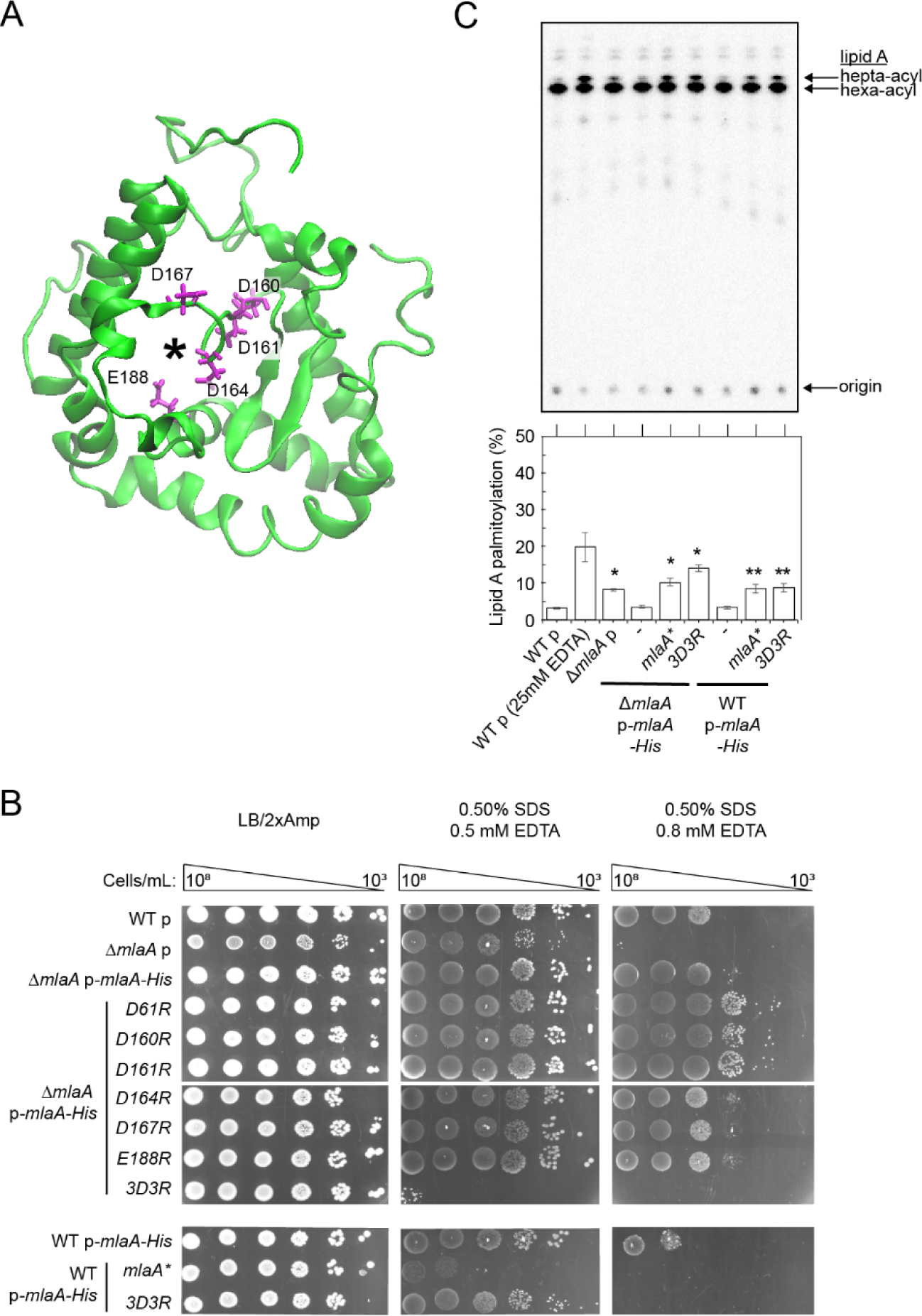
A triple charge-inversion mutation in the hydrophilic channel within MlaA results in gain-of-function phenotypes. (*A*) A representative structure of MlaA from all-atomistic MD simulations (as in Fig. 3, *top view*) with position of the channel depicted by an *asterisk*. Negatively charged residues mutated to arginine are highlighted (*magenta*, in sticks). (*B*) Analysis of SDS/EDTA sensitivity of wild-type (WT) and Δ*mlaA* strains producing indicated MlaA variants at low levels from the pET23/42 vector (p) (12). Serial dilutions of respective cultures were spotted on LB agar plates containing Amp, supplemented with or without 0.50% SDS and 0.5/0.8 mM EDTA, as indicated, and incubated overnight at 37 °C. (*C*) Representative thin layer chromatography (TLC)/autoradiographic analysis of [^32^P]-labeled lipid A extracted from exponential phase cultures of strains described in (*B*). As a positive control for lipid A palmitoylation, WT cells were treated with 25 mM EDTA for 10 min prior to extraction. Equal amounts of radioactive material were spotted for each sample. Average percentages of palmitoylation of lipid A and the standard deviations were quantified from triplicate experiments and plotted below. Student’s t-tests: *, p < 0.005 compared to WT with empty vector; **, p < 0.001 compared to WT p-*mlaA-His*.

### Flexibility of a hairpin loop adjacent to the channel on MlaA is required for function

The gain-of-function/dominant negative phenotype of the *mlaA_3D3R_* mutant is similar to a previously reported *mlaA\** (or *mlaA_ΔNF_*) mutant (Figs. 5*B* and 5*C*) (23), suggesting that these mutations may have similar effects on MlaA structure and/or function. Interestingly, the positions of these mutations on the OmpC-MlaA models flank a hydrophobic hairpin loop (G141-L158) within MlaA (Fig. 6*A*). Therefore, we hypothesized that the loop could play a functional role in MlaA, and that these mutations may affect interactions with this loop. To examine this possibility, we created three separate mutations at the hairpin structure and tested each variant for MlaA function. Two of these mutations, Y^147^VQL→4A (*L1*) and F^152^YGSF→5A (*L2*), are designed to disrupt interactions with other regions of MlaA. The other mutation, P151A, removes a proline that may be critical for the hairpin turn structure. The N-terminus of the loop is connected to the rest of MlaA via an unstructured glycine-rich linker, which we reasoned may influence conformation of the entire hairpin structure. Thus, we constructed two additional mutants, G^141^VGYG→A^141^VAYA (*3G3A*) and G^141^VGYG→P^141^VPYP (*3G3P*), to reduce possible flexibility in this region. Remarkably, *L1*, *L2*, and *3G3P* mutations resulted in similar extents of SDS/EDTA sensitivity (Fig. 6*B*), as well as OM outer leaflet PL accumulation (Fig. 6*C*), when compared to the Δ*mlaA* mutation. Given that these mutations also do not affect MlaA levels or interaction with OmpC (Fig. S11), we conclude that they are loss-of-function mutations. The hairpin loop, along with its surrounding structures, forms an important functional region on MlaA.

**Figure 6.**
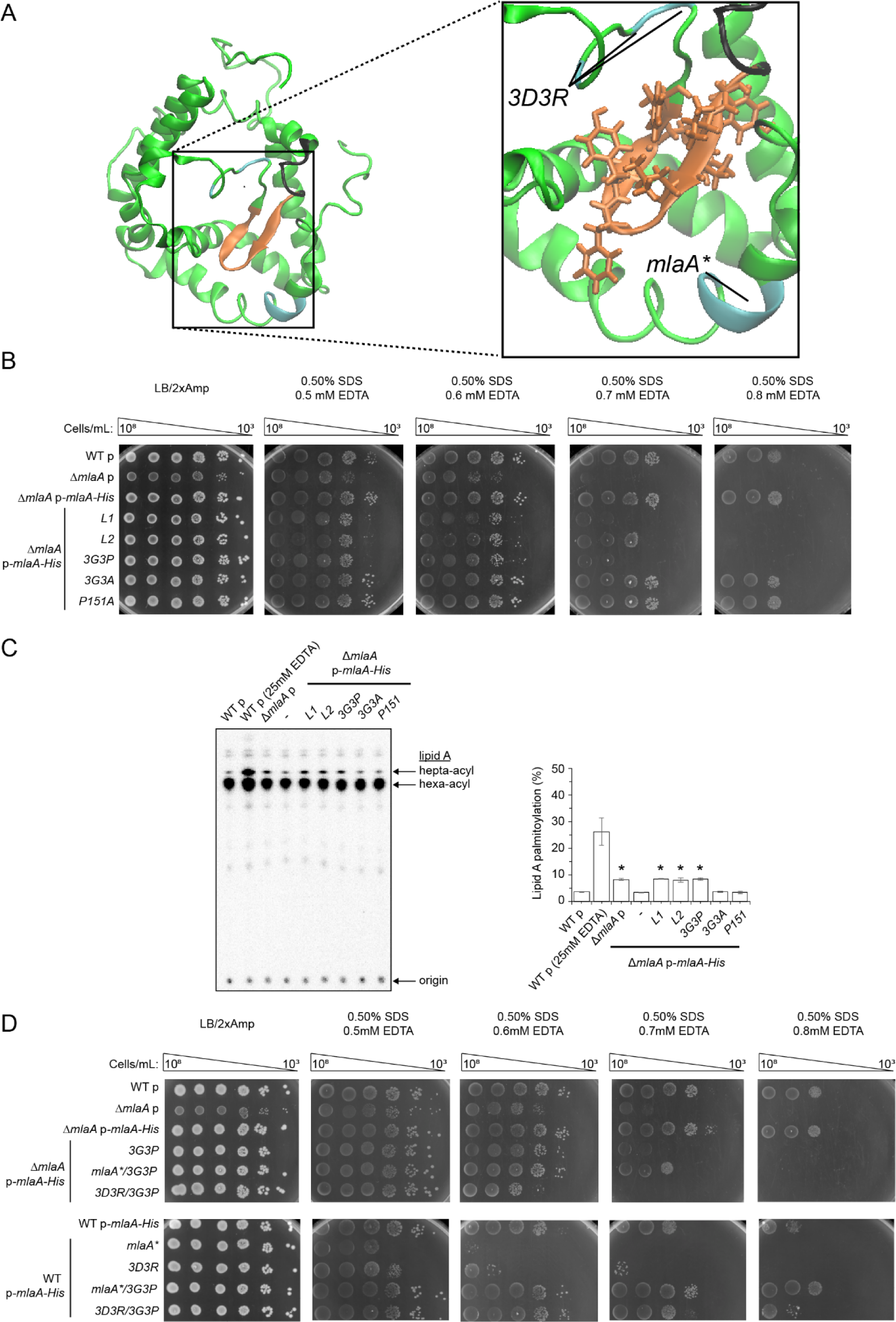
Flexibility in a hairpin loop structure on MlaA adjacent to the hydrophilic channel is critical for function. (*A*) A representative structure of MlaA from all-atomistic MD simulations (as in Fig. 3*B*) with the hairpin loop adjacent to the hydrophilic channel highlighted. In the expanded representation, the *3D3R* and *mlaA\** mutations, the hairpin loop, and the glycine rich region N-terminal to the loop are colored in *cyan*, *orange* and *black*, respectively. Residues on the hairpin loop chosen for mutation are represented in sticks. (*B*) Analysis of SDS/EDTA sensitivity of wild-type (WT) and Δ*mlaA* strains producing indicated MlaA loop variants from the pET23/42 vector (p). (*C*) Representative TLC/autoradiographic analysis of [^32^P]-labeled lipid A extracted from exponential phase cultures of strains described in (*B*). Equal amounts of radioactive material were spotted for each sample. Average percentages of palmitoylation of lipid A and the standard deviations were quantified from triplicate experiments and plotted on the right. Student’s t-tests: *, p < 0.0005 compared to WT with empty vector. (*D*) Analysis of SDS/EDTA sensitivity of wild-type (WT) and Δ*mlaA* strains producing indicated MlaA variants from the pET23/42 vector (p).

Phenotypes observed for the loop rigidifying mutation (*3G3P*) and gain-of-function mutations (*3D3R* and *mlaA\**) suggest that flexibility in the hairpin loop is critical for MlaA function. We hypothesize that the hairpin loop may exist in two distinct conformations. The *3D3R* or *mlaA\** mutations could alter interactions with the loop, resulting in it adopting one conformation, and somehow giving rise to gain-of-function/dominant negative phenotypes; in the case of *mlaA\**, it was proposed that this mutation caused MlaA to be in a “leaky” or “open” state, and allowed PLs to flip out to the outer leaflet of the OM (23). In contrast, the *3G3P* mutation may lock the hairpin loop in a second conformation, where MlaA is in a “closed” state, thus abolishing function in PL transport. If these were true, we predict that rigidification of the hairpin loop with the *3G3P* mutation would be able to correct gain-of-function/dominant negative phenotypes observed in the *3D3R* and/or *mlaA\** mutants. Indeed, the *3G3P/3D3R* and *3G3P/mlaA\** combination mutants no longer exhibit gain-of-function/dominant negative phenotypes, but behave like the *3G3P* or null mutants (Fig. 6*D*). Again, these variants are expressed at comparable levels to the single mutants, and still interact with OmpC (Fig. S11). Taken together, these results indicate clear importance of dynamics in the hairpin loop in controlling the function of MlaA.

### A specific residue in the dimeric interface of OmpC is important for its function in maintaining lipid asymmetry

Given that MlaA sits in the membrane and provides a channel that putatively allows movement of PLs across the OM, it is not clear why it should bind at one of the dimeric interfaces of OmpC trimers, and what the exact role of OmpC may be. To understand the importance of OmpC-MlaA interaction, we attempted to engineer monomeric OmpC constructs that we predict would no longer interact with MlaA. We installed specific mutations (G19W and/or R92L) in OmpC that were found previously to disrupt the oligomerization state of its homolog OmpF (G19W and R100L correspondingly) in vitro (24) (Fig. 7*A*). Both the OmpC_G19W_ and OmpC_R92L_ single mutants can interact with MlaA, and still form trimers in vitro, albeit slightly destabilized compared to wild-type OmpC (Fig. S12). Combination of these mutations further weakens the OmpC trimer, with noticeable monomer population at physiological temperature. Intriguingly, both the double mutant and the R92L single mutant accumulated PLs in the outer leaflet of the OM (Fig. 7*B*), indicating that R92 is important for the role of OmpC in OM lipid asymmetry. Consistent with this idea, we demonstrated that cells expressing the OmpC_R92A_ variant also exhibit perturbations in OM lipid asymmetry. We further showed that all these *ompC* alleles can rescue severe SDS/EDTA sensitivity known for cells lacking OmpC (Fig. 7*C*), suggesting normal porin function; however, cells expressing OmpCG19W/R92L and OmpC_R92A_, unlike WT and OmpC_R92L_, are still sensitive to SDS at higher concentrations of EDTA. These phenotypes mirror those observed for cells lacking MlaA, suggesting the loss of Mla function in these mutant strains. It appears that the R92 residue is critical for this function, although it is not clear why the single R92L mutation did not result in SDS/EDTA sensitivity. The exact role of R92 is not known, but the phenotypes observed for the single R92A mutation cannot be due to disruption of OmpC trimerization (Fig. S12). We conclude that OmpC has an active role in maintaining OM lipid symmetry together with MlaA.

**Figure 7.**
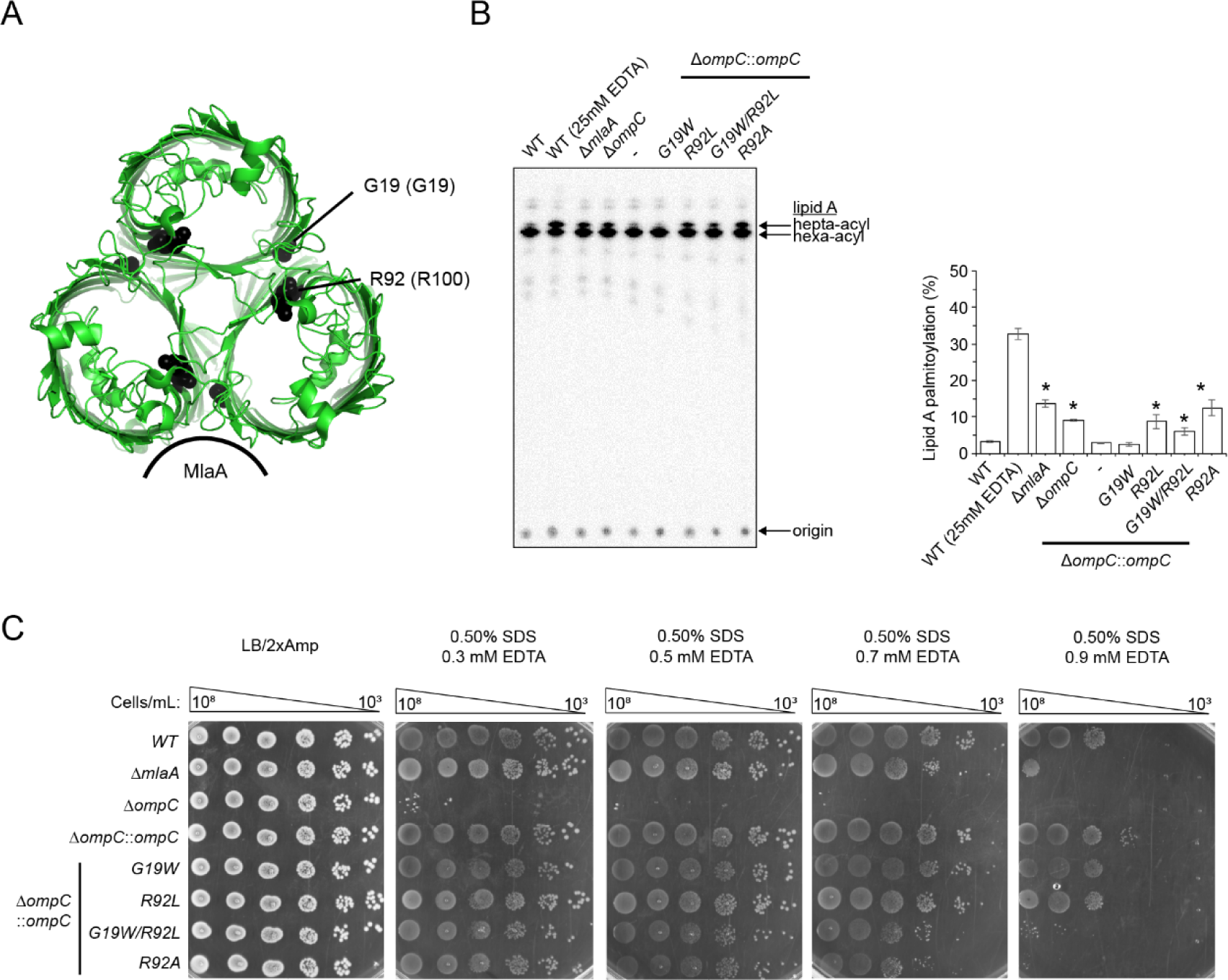
A specific mutation in the dimeric interface of the OmpC trimer results in perturbation in OM lipid asymmetry. (*A*) Cartoon representation of the crystal structure of OmpC trimer illustrating the positions of G19 and R92 region. (*B*) Analysis of SDS/EDTA sensitivity of wild-type (WT) and Δ*ompC* strains producing indicated OmpC variants from the chromosomal locus. (*C*) Representative TLC/autoradiographic analysis of [^32^P]-labeled lipid A extracted from stationary phase cultures of strains described in (*B*). Equal amounts of radioactive material were spotted for each sample. Average percentages of palmitoylation of lipid A and the standard deviations were quantified from triplicate experiments and plotted on the right. Student’s t-tests: *, p < 0.005 compared to Δ*ompC*::*ompC*.

## Discussion

Osmoporin OmpC interacts with MlaA to maintain lipid asymmetry in the OM; how this complex is organized to extract PLs from the outer leaflet of the OM is not known. In this study, we have employed photoactivatable crosslinking and MD simulations to gain insights into the molecular architecture of the OmpC-MlaA complex. We have established that MlaA interacts extensively with OmpC at one of the dimeric interfaces of the porin trimer, and resides entirely within the OM lipid bilayer. We have also demonstrated that MlaA forms a hydrophilic channel, likely allowing PLs to translocate across the OM. This overall organization of the OmpC-MlaA complex is quite remarkable, especially how MlaA spans the OM and possibly gains access to outer leaflet PLs. Very few lipoproteins are known to span the OM; some notable examples include the LptDE and Wza translocons, which transports LPS and capsular polysaccharides, respectively. In the LptDE complex, the OM lipoprotein LptE serves as a plug, and stretches across the bilayer through the lumen of the LptD β-barrel (25, 26). In the octameric Wza translocon, each protomer provides a C-terminal α-helix to form a pore that spans the membrane (27). MlaA is unique in that it is essentially an integral membrane protein, capable of forming a channel on its own. In many ways, MlaA behaves like typical OM β-barrel proteins, even though it is predominantly α-helical. Furthermore, being overall a hydrophobic protein also poses a problem for MlaA to transit across the periplasmic space. How MlaA is shielded from the aqueous environment, in addition to the requirement of the Lol system (28), necessitates further investigation.

Very recently, crystal structures of MlaA in complex with trimeric porins have in fact been solved (29). Interestingly, the experimentally-determined structures of MlaA closely resembled the initial MlaA model predicted from co-evolution analysis (Fig. S13*A*). These structures revealed that MlaA interacts with trimeric porins at one or more of their dimeric interfaces (Fig. S13*B*), in an orientation similar to one of our simulated OmpC-MlaA models (Fig. 3*B*). They also showed that MlaA contains a channel. Clearly, these findings converged with the main conclusions derived from our biochemical and modelling studies, which is quite remarkable. We note, however, a couple of discrepancies between the solved structures and our biochemical data, suggesting that these static structures may represent only one of several possible stable conformations of MlaA, which may exist as part of the mechanism associated with lipid transport in the native OM environment. First, the distance separating porin residues equivalent to Y149/L340 in OmpC and the MlaAF133-R205 peptide (Fig. S13*B*) is not consistent with the detection of strong photoactivatable crosslinks between these regions in the complex (Fig. 2*B*). Second, the proposed location of some MlaA residues, specifically M39, F42, and N43, at the interior of the lipid bilayer (Fig. S13*B*) is not in agreement with high or partial solvent accessibility of these sites in cells, as highlighted in the SCAM data (Fig. S9*B*). Upon close examination of the structures, we found multiple non-native MlaA-MlaA and MlaA-porin contacts in all the crystal forms (Fig. S13*C*); these artificial crystal contacts, some with substantial buried surface areas, may have influenced the observed conformation of MlaA. Given that the MlaA-porin structures were also not solved in the context of a native lipid bilayer, we suspect that they do not yet provide the complete picture.

Our functional data on the OmpC-MlaA complex provide a glimpse of how PLs may translocate across the OM during maintenance of lipid asymmetry. One key aspect of MlaA function resides in a hairpin loop structure juxtaposed against the hydrophilic channel. Dynamics of this loop appear to control whether MlaA exists in a “closed” or “open” state, and thus access of PLs through the channel. A mutation that likely rigidifies the loop locks MlaA in the nonfunctional “closed” state (Fig. 6), while mutations that possibly affect interactions with the loop favors the “open” state, and gives rise to gain of function (Fig. 5). Therefore, the hairpin loop may directly gate the channel, providing access for outer leaflet PLs across the OM. This idea has also been suggested based on the solved MlaA structures, where a putative disulfide bond that apparently locks the loop in position renders MlaA non-functional (29).

How OmpC participates in maintaining OM lipid asymmetry as part of the complex is not clear, especially given that MlaA alone provides a channel for PL translocation. It is possible that OmpC may play a passive role, and simply be important for stabilizing the structure and orientation of MlaA in the OM. However, we have previously shown that MlaA also interacts with OmpF, yet removing OmpF has minimal effects on OM lipid asymmetry (12); this argues against a mere passive role for OmpC in PL translocation. Furthermore, we have now identified a specific residue R92 on OmpC that is required for maintaining OM lipid asymmetry (Fig. 7). Therefore, we believe that OmpC plays an active role in the process. R92 lies in the dimeric interface of the OmpC trimer, which incidentally is where MlaA binds. Even though this residue has been shown to be important for gating the porin (30), it is not obvious how this gating function may influence the translocation of PLs by MlaA. Instead, we speculate thatbeing close to Y149 and L340, R92 may somehow affect interactions between OmpC and the MlaAF133-R205 peptide, possibly influencing MlaA conformation in the OM, and in turn, properties of the hydrophilic channel. The role that R92 plays in OmpC-MlaA function should be further characterized.

The OmpC-MlaA complex is proposed to extract PLs from the outer leaflet of the OM and hand them over to MlaC, which resides in the periplasm. Consistent with this idea, *E. coli* MlaC has been crystallized with a bound PL, and shown to interact with a complex of OmpF-MlaA in vitro (14). However, it is not clear how transfer of PLs from OmpC-MlaA to MlaC takes place, even though this must presumably occur in an energy-independent fashion. Since PL movement from the outer to inner leaflets of the OM is entropically disfavored, it is likely that translocation of PLs by the OmpC-MlaA complex would be coupled to transfer to MlaC, i.e. extracted PLs do not go into the inner leaflet. If this were true, it may be possible that MlaC also influences the function of the OmpC-MlaA complex. Specifically, binding of MlaC to the complex may be required for PL extraction from the outer leaflet of the OM. MlaC could alter the structure and/or dynamics of MlaA in the OmpC-MlaA complex, ultimately leading to efficient PL translocation across the OM.

Lipid asymmetry is critical for the OM to function as an effective permeability barrier. Thus, understanding mechanistic aspects of how bacterial cells maintain OM lipid asymmetry would guide us in designing strategies to overcome the barrier. Our work on elucidating the architecture and function of the OmpC-MlaA complex has revealed critical insights into the role of a hairpin loop on MlaA in modulating activity, a feature that can be exploited in drug discovery efforts. In particular, small molecules that can potentially influence dynamics of this loop may induce either loss or gain of function, thereby leading to increased sensitivity to existing antibiotics.

## Experimental procedures

### Bacterial strains and plasmids

All strains and plasmids used are listed in Table S1 and S2, respectively.

### Growth conditions

Luria Bertani (LB) broth and agar were prepared as described previously (12). Unless otherwise noted, ampicillin (Amp) (Sigma-Aldrich, MO, USA) was used at a concentration of 200 μg/mL, chloramphenicol (Cam) (Alfa Aesar, Heysham, UK) at 15 μg/mL, kanamycin (Kan) (Sigma-Aldrich) at 25 μg/mL, and spectinomycin (Spec) (Sigma-Aldrich) at 50 μg/mL. For crosslinking experiments, *para*-Benzoyl-L-phenylalanine (*p*Bpa; Alfa Aesar) was dissolved in 1 M NaOH at 0.25 M, and used at 0.25 mM unless otherwise mentioned.

### Plasmid construction

To construct most plasmids, the desired gene or DNA fragments were amplified by PCR from the DNA template using primers listed in Table S3. Amplified fragments were digested with relevant restriction enzymes (New England Biolabs) and ligated into the same sites of an appropriate plasmids using T4 DNA ligase (New England Biolabs). NovaBlue competent cells were transformed with the ligation products and selected on LB plates containing appropriate antibiotics. All constructs were verified by DNA sequencing (Axil Scientific, Singapore).

### Construction of chromosomal *ompC* mutants using negative selection

All chromosomal *ompC* mutations were introduced via a positive-negative selection method described previously (31). To prepare electro-competent cells, strain MC4100 harbouring pKM208 (32) grown overnight at 30 °C was inoculated into 15 mL SOB broth with 1:100 dilution. Cells were grown at the same temperature until OD_600_ reached ˜0.3-0.4. 1 mM of IPTG was added and the culture was grown for another 60 min at 30 °C. Cells were then subjected to heat shock at 42 °C for 15 min, followed by incubation for 15 min on ice, with intermittent agitation. Subsequently, cells were centrifuged at 5000 x *g* for 10 min and made competent by washing twice with prechilled sterile water followed by cold 10 % glycerol. Competent cells were pelleted and resuspended in cold 10 % glycerol. For positive selection, 1 μg *kan-PrhaB-tse2* cassette amplified from pSLC-246 (31) using ompC_NS_N5 and ompC_NS_C3 primer pairs was transformed into the competent cells using 1-mm electroporation cuvettes (Biorad) in Eppendorf Eporator^®^ (Eppendorf) with an output voltage of 1800 V. Cells were recovered in LB with 2 % glucose at 37 °C for at least 4 h, plated onto LB plates supplemented with Kan and 2 % glucose, and incubated at 37 °C for 24 h. For negative selection, 1 μg PCR product of *ompC* wild-type or mutant constructs amplified using ompC_NS_N5_C and ompC_NS_C3_C primer pairs were transformed into competent cells made from positive selection using similar procedures. After transformation, cells were plated onto minimal (M9) plates supplemented with 0.2 % rhamnose, and incubated for 48 h at 37 °C. Surviving colonies were PCR screened and verified by DNA sequencing (Axil Scientific, Singapore).

### In vivo photoactivatable crosslinking

We adopted previously described protocol (19) for all in vivo photoactivable crosslinking experiments. Briefly, amber stop codon (TAG) was introduced at selected positions in either pDSW206*ompC* or pCDF*mlaA-His* plasmids via site directed mutagenesis using primers listed in Table S3. For OmpC crosslinking, MC4100 with Δ*ompC*::*kan* background harbouring p*Sup-BpaRS-6TRN* (33) and pDSW206*ompC* were used. For MlaA crosslinking, MC4100 with Δ*mlaA*::*kan* background harbouring p*Sup-BpaRS-6TRN* (33) and pCDF*mlaA-His* were used. An overnight 5 mL culture was grown from a single colony in LB broth supplemented with appropriate antibiotics at 37 °C. Overnight cultures were diluted 1:100 into 10 mL of the same media containing 0.25 mM *p*Bpa and grown until OD_600_ reached ˜1.0. Cells were normalized by optical density before pelleting and resuspended in 1 mL ice cold TBS (20 mM Tris pH 8.0, 150 mM NaCl). Samples were either used directly or irradiated with UV light at 365 nm for 20 min at 4 °C or room temperature. All samples were pelleted again and finally resuspended in 200 μL of 2 X Laemmli buffer, boiled for 10 min, and centrifuged at 21,000 x *g* in a microcentrifuge for one min at room temperature; 15 μL of each sample subjected to SDS-PAGE and immunoblot analyses.

### Over-expression and purification of OmpC-MlaA-His complexes

All proteins were overexpressed in and purified from BL21(λDE3) derivatives. We found that BL21(λDE3) strains from multiple labs do not actually produce OmpC (gene is somehow missing in these strains); therefore, to obtain OmpC-MlaA complexes, we deleted *ompF* from the chromosome and introduced *ompC* on a plasmid. OmpC-MlaA-His protein complexes were over-expressed and purified from BL21(λDE3) cells with chromosomal Δ*ompF*::*kan* background co-transformed with either pDSW206*ompC_p_*_Bpa_, p*Sup-BpaRS-6TRN* and pCDF*dmlaA-His* (for in vitro crosslinking experiments), or pACYC184*ompC* and pET22b(+)*dmlaA-His* (for characterization of the wildtype complex). An overnight 10 mL culture was grown from a single colony in LB broth supplemented with appropriate antibiotics at 37 °C. The cell culture was then used to inoculate a 1-L culture and grown at the same temperature until OD_600_ reached ˜ 0.6. For induction, 0.5 mM IPTG (Axil Scientific, Singapore) was added and the culture was grown for another 3 h at 37 °C. Cells were pelleted by centrifugation at 4,700 x *g* for 20 min and then resuspended in 10-mL TBS containing 1 mM PMSF (Calbiochem) and 30 mM imidazole (Sigma-Aldrich). Cells were lysed with three rounds of sonication on ice (38 % power, 1 second pulse on, 1 second pulse off for 3 min). Cell lysates were incubated overnight with 1 % ndodecyl β-D-maltoside (DDM, Calbiochem) at 4 °C. Cell debris was removed by centrifugation at 24,000 x *g* for 30 min at 4 °C. Subsequently, supernatant was incubated with 1 mL Ni-NTA nickel resin (QIAGEN), pre-equilibrated with 20 mL of wash buffer (TBS containing 0.025 % DDM and 80 mM imidazole) in a column for 1 h at 4 °C with rocking. The mixture was allowed to drain by gravity before washing vigorously with 10 x 10 mL of wash buffer and eluted with 10 mL of elution buffer (TBS containing 0.025% DDM and 500 mM imidazole). The eluate was concentrated in an Amicon Ultra 100 kDa cut-off ultra-filtration device (Merck Millipore) by centrifugation at 4,000 x *g* to ˜500 μL. Proteins were further purified by SEC system (AKTA Pure, GE Healthcare, UK) at 4 °C on a prepacked Superdex 200 increase 10/300 GL column, using TBS containing 0.025% DDM as the eluent. Protein samples were used either directly or irradiated with UV at 365 nm for in vivo photoactivable crosslinking experiments.

### SEC-MALS analysis to determine absolute molar masses of OmpC-MlaA-His complex

Prior to each SEC-MALS analysis, a preparative SEC was performed for BSA (Sigma-Aldrich) to separate monodisperse monomeric peak and to use as a quality control for the MALS detectors. In each experiment, monomeric BSA was injected before the protein of interest and the settings (calibration constant for TREOS detector, Wyatt Technology) that gave the well-characterized molar mass of BSA (66.4 kDa) were used for the molar mass calculation of the protein of interest. SEC purified OmpC-MlaA-His was concentrated to 5 mg/mL and injected into Superdex 200 Increase 10/300 GL column pre-equilibrated with TBS and 0.025 % DDM. Light scattering (LS) and refractive index (*n*) data were collected online using miniDAWN TREOS (Wyatt Technology, CA, USA) and Optilab T-rEX (Wyatt Technology, CA, USA), respectively, and analyzed by ASTRA 6.1.5.22 software (Wyatt Technology). Protein-conjugate analysis available in ASTRA software was applied to calculate non-proteinaceous part of the complex. In this analysis, the refractive index increment *dn/dc* values (where *c* is sample concentration) of 0.143 mL/*g* and 0.185 mL/*g* were used for DDM and protein complex, respectively (34). For BSA, UV extinction coefficient of 0.66 mL/(mg.cm) was used. For the OmpC-MlaA-His complex, that was calculated to be 1.66 mL/(mg.cm), based on its predicted stoichiometric ratio OmpC_3_MlaA.

### Affinity purification experiments

Affinity purification experiments were conducted using Δ*mlaA* strains expressing MlaA-His at low levels from the pET23/42 vector. For each strain, a 1.5-L culture (inoculated from an overnight culture at 1:100 dilution) was grown in LB broth at 37 °C until OD_600_ of ˜0.6. Cells were pelleted by centrifugation at 4700 x *g* for 20 min and then resuspended in 10-mL TBS containing 1 mM PMSF (Calbiochem) and 50 mM imidazole (Sigma-Aldrich). Cells were lysed with three rounds of sonication on ice (38 % power, 1 second pulse on, 1 second pulse off for 3 min). Cell lysates were incubated overnight with 1 % ndodecyl β-D-maltoside (DDM, Calbiochem) at 4 °C. Cell debris was removed by centrifugation at 24,000 x *g* for 30 min at 4 °C. Subsequently, supernatant was incubated with 1 mL Ni-NTA nickel resin (QIAGEN), pre-equilibrated with 20 mL of wash buffer (TBS containing 0.025% DDM and 80 mM imidazole) in a column for 1 h at 4 °C with rocking. The mixture was allowed to drain by gravity before washing vigorously with 10 x 10 mL of wash buffer and eluted with 5 mL of elution buffer (TBS containing 0.025% DDM and 500 mM imidazole). The eluate was concentrated in an Amicon Ultra 100 kDa cut-off ultra-filtration device (Merck Millipore) by centrifugation at 4,000 x *g* to ˜100 μL. The concentrated sample was mixed with equal amounts of 2X Laemmli buffer, boiled at 100 °C for 10 min, and subjected to SDS-PAGE and immunoblot analyses.

### Trypsin digestion for protein N-terminal sequencing and mass spectrometry analyses

A 1 mg/mL solution of purified OmpC-MlaA-His (OmpC was either wild-type or substituted with *p*Bpa at selected positions) complex was incubated with or without 50 μg/mL trypsin (Sigma-Aldrich) for 1 h at room temperature. *p*Bpa substituted samples were irradiated with UV at 365 nm before trypsin digestion. Samples were then analyzed by SDS-PAGE, followed by SEC. Peak fractions from SEC for each sample were pooled, concentrated using an Amicon Ultra 100 kDa cut-off ultra-filtration device (Merck Millipore), and resuspended in 2 X Laemmli sample buffer before analyses by SDS-PAGE and immunoblot using α-MlaA antibody. For N-terminal sequencing, samples were transferred onto PVDF membrane, followed by Coomassie Blue staining (1–2 s). The desired protein bands were carefully excised with a surgical scalpel. For tandem MS, protein bands were excised from a Coomassie Blue stained Tricine gel. Samples prepared for N-terminal sequencing and tandem MS were kept in sterile 1.5 mL centrifuge tubes before submission for analyses at Tufts University Core Facility, Boston, USA, and Taplin Biological Mass Spectrometry Facility, Harvard Medical School, Boston, USA, respectively.

### Substituted cysteine accessibility method (SCAM)

1-mL cells were grown to exponential phase (OD_600_ ˜0.6), washed twice with TBS (pH 8.0), and resuspended in 480 μL of TBS. For the blocking step, four tubes containing 120 μL of cell suspension were either untreated (positive and negative control tubes added with deionized H_2_O) or treated with 5 mM thiol-reactive reagent *N*-ethylmaleimide (NEM, Thermo Scientific) or sodium (2-sulfonatoethyl) methanethiosulfonate (MTSES, Biotium). As MTSES is membrane impermeable, it is expected to react with the free cysteine in MlaA variants only when the residue near or at the membrane-water boundaries, or in a hydrophilic channel. In contrast, NEM is expected to label all MlaA cysteine variants as it is membrane permeable. Reaction with MTSES or NEM blocks the particular cysteine site from subsequently labelling by maleimide-polyethylene glycol (MalPEG; 5 kDa, Sigma-Aldrich). After agitation at room temperature for 1 h, cells were washed twice with TBS, pelleted at 16,000 x *g*, and resuspended in 100 μL of lysis buffer (10 M urea, 1% SDS, 2 mM EDTA in 1 M Tris pH 6.8). Both NEM- and MTSES-blocked samples and the positive control sample were exposed to 1.2 mM Mal-PEG-5k. After agitation for another hour with protection from light, all samples were added with 120 μL of 2 X Laemmli buffer, boiled for 10 min, and centrifuged at 21,000 x *g* in a microcentrifuge for one min at room temperature; 20 μL from each sample tubes were subjected to SDS-PAGE and immunoblot analyses.

### In vivo disulfide bond analysis

Strain NR1216 (Δ*dsbA*) harbouring pET23/42*mlaA-His* expressing MlaA-His with site specific cysteine substitutions was grown overnight in LB broth at 37 °C. A 0.5 mL of overnight culture was normalized by optical density, added with trichloroacetic acid (TCA) at final concentration of ˜14 % and mixed thoroughly at 4 °C. This step was performed to prevent scrambling of disulfide bond formed in the cysteines substituted MlaA-His. Proteins precipitated for at least 30 min on ice were centrifuged at 16,000 × *g* for 10 min at 4 °C. The pellet was washed with 1 mL of ice-cold acetone and centrifuged again at 16,000 × *g* for 10 min at 4 °C. Supernatants were then aspirated and the pellet was air dried at room temperature for at least 20 min. Samples were resuspended thoroughly with 100 μL of either 100 mM Tris.HCl pH 8.0, 1% SDS (for non-reduced samples), or the same buffer supplemented with 100 mM of dithiothreitol (DTT) (for reduced samples), incubated for 20 min at room temperature. The samples were finally mixed with 4 X Laemmlli buffer, boiled for 10 min and subjected to SDS-PAGE and immunoblotting analyses using α-His antibody.

### Docking of MlaA to OmpC

The ClusPro server (22) was used to dock MlaA (ligand, uniprot ID: P76506, https://gremlin2.bakerlab.org/meta_struct.php?page=p76506) (17) to OmpC (receptor, PDB ID: 2J1N) (18). The default server settings were used in the docking procedure. The minimum distance between 6 residues on OmpC and the corresponding cross-linked peptide regions of MlaA was calculated for all the predicted structures obtained from the server. Two OmpC-MlaA model with the smallest average minimum distance of all residue and peptide pairs were selected as the initial structures for use in the all-atom simulations.

### Simulation procedures and setup

All simulations were performed using version 5.1.4 of the GROMACS simulation package (35, 36).

*All-atom simulations*. In total, 7 all-atom simulations were performed (Table A). The simulations were performed using the CHARMM36 force filed parameter set (37). The equations of motion were integrated using the Verlet leapfrog algorithm with a step size of 2 fs. Lengths of hydrogen bonds were constrained with the LINCS algorithm (38). Electrostatic interactions were treated using the smooth Particle Mesh Ewald (PME) method (39), with cutoff for short-range interactions of 1.2 nm. The van der Waals interactions were switched smoothly to zero between 1.0 and 1.2 nm. The neighbor list was updated every 20 steps. The Nose-Hoover thermostat (40, 41) with a coupling constant of 1.0 ps was used to maintain a constant system temperature of 313 K. The protein, membrane and solvent (water and ions) were coupled to separate thermostats. The Parrinello-Rahman barostat (42) with a coupling constant of 5.0 ps was used to maintain a pressure of 1 bar. Semi-isotropic pressure coupling was used for all the membrane systems, while isotropic coupling was used for the solvent-only system. Initial velocities were set according to the Maxwell distribution.

Proteins were inserted into a pre-equilibrated, symmetrical 1,2-dimyristoylphosphatidylethanolamine (DMPE) membrane over 5 ns using the membed tool (43) in the GROMACS simulation package. Subsequent equilibration, with position restraints of 1000 kJ mol^-1^ placed on all non-hydrogen protein atoms, was performed for 20 ns to allow the solvent and lipids to equilibrate around the proteins. The position restraints were removed before performing the production runs.

**Table A.** Summary of all-atom molecular simulations: system compositions and simulation times.

| Protein Configuration | Lipids | Water and Ions | Simulation time (# of simulations x ns) |
| --- | --- | --- | --- |
| MlaA | N/A | 9439 H<sub>2</sub>O<br>29 K<sup>+</sup><br>19 Cl<sup>-</sup> | 1 x 500 |
| MlaA | 272 DMPE | 11734 H<sub>2</sub>O<br>42 K<sup>+</sup><br>32 Cl<sup>-</sup> | 1 x 500 |
| OmpC trimer<br>MlaA (ClusPro model) in orientation 1 | 980 DMPE | 36113 H<sub>2</sub>O<br>98 K<sup>+</sup><br>98 Cl<sup>-</sup> | 1 x 500<br>1 x 320<br>1 x 130 |
| OmpC trimer<br>MlaA (ClusPro model) in orientation 2 | 980 DMPE | 36113 H<sub>2</sub>O<br>98 K<sup>+</sup><br>98 Cl<sup>-</sup> | 1 x 500<br>1 x 500 |

All of the simulations performed are summarized in Table A. For the OmpC trimer docked with MlaA in orientation 1, three separate production simulations with different initial velocities were performed for the OmpC-MlaA complex, resulting in 3 trajectories of 500 ns, 320 ns, and 130 ns in length, respectively. For the OmpC trimer docked with MlaA in orientation 2, two separate production simulations with different initial velocities were performed for the complex, resulting in 2 trajectories of 500 ns in length. Clustering was performed on the MlaA structures obtained from a combined trajectory of all three (MlaA orientation 1) or two (MlaA orientation 2) atomistic simulations – a total of 4750 frames spaced every 0.2 ns. The structures were assigned to clusters using Root Mean Squared Distance (RMSD) with a 0.1 nm cut-off. Four and two clusters, respectively, were observed for MlaA in the two orientations to contain greater than 100 frames. The central structure of these four clusters was used to generate the representative OmpC-MlaA models (Fig. S7). trj_cavity was used to identify the location the pore cavity (44), and Hole was used to create the pore profile (45).

### Temperature titration for chromosomal *ompC* mutants

Purified wild-type and mutant OmpC-MlaA-His complexes were aliquoted into 1.5 mL centrifuge tubes and incubated in water bath set at different temperatures for 10 min. 20 μl of each sample were transferred into separate tubes and mixed immediately with equal volume of 2 X Laemmlli buffer and subjected to SDS-PAGE in 12 % Tris.HCl gels, followed by Coomassie Blue staining (Sigma-Aldrich).

### LPS labeling and lipid A isolation

Mild acid hydrolysis of [^32^P]-labeled cultures was used to isolate lipid A according to a procedure described previously (4, 12, 52) with some modifications. 5 mL cultures were grown (inoculated with overnight cultures at 1:100 dilution) in LB broth at 37 °C until OD_600_ reached ˜0.5–0.7 (exponential) or ˜2–4 (stationary). Cultures were uniformly labeled with 1 μCi mL–1 [^32^P]-disodium phosphate (Perkin-Elmer) from the start of inoculation. One MC4100 wild-type culture labeled with [^32^P] was treated with 25 mM EDTA, pH 8.0 for 10 min prior to harvesting. Cells (5 mL and 1 mL for exponential and stationary phase cultures respectively) were harvested by centrifugation at 4700 × *g* for 10 min and washed twice with 1 mL PBS (137 mM NaCl, 2.7 mM KCl, 10 mM Na_2_HPO_4_, 1.8 mM KH_2_PO_4_, pH 7.4) at 5000 × *g* for 10 min. Each cell pellet was resuspended in 0.32 mL PBS, and converted into single phase Bligh/Dyer mixture (chloroform/methanol/water:1/2/0.8) by adding 0.8 mL methanol and 0.4 mL chloroform. The single phase Bligh/Dyer mixture was incubated at room temperature for 20 min, followed by centrifugation at 21,000 × *g* for 30 min. Each pellet obtained was washed once with 1 mL freshly made single phase Bligh/Dyer mixture and centrifuged as above. The pellet was later resuspended in 0.45 mL 12.5 mM sodium acetate containing 1 % SDS, pH 4.5. The mixture was sonicated for 15 min before incubation at 100 °C for 40 min. The mixture was converted to a two-phase Bligh/Dyer mixture (chloroform/methanol/water: 2/2/1.8) by adding 0.50 mL methanol and 0.50 mL chloroform. The lower phase of each mixture was collected after phase partitioning by centrifugation at 21 000 × *g* for 30 min. The collected lower phase was washed once with 1 mL of the upper phase derived from the freshly made two-phase Bligh/Dyer mixture and centrifuged as above. The final lower phase was collected after phase partitioning by centrifugation and dried under N_2_ gas. The dried radiolabeled lipid A samples were redissolved in 100 μL of chloroform/methanol mixture (4/1), and 20 μL of the samples were used for scintillation counting (MicroBeta2®, Perkin-Elmer). Equal amounts of radiolabeled lipids (cpm/lane) were spotted onto the TLC plate (Silica Gel 60 F254, Merck Millipore) and were separated using the solvent system consisting of chloroform/pyridine/96 % formic acid/water (50/50/14.6/4.6) (5). The TLC plate was then dried and exposed to phosphor storage screens (GE Healthcare). Phosphor-screens were visualized in a phosphor-imager (Storm 860, GE Healthcare), and the spots were analyzed by ImageQuant TL analysis software (version 7.0, GE Healthcare). Spots were quantified and averaged based on three independent experiments of lipid A isolation.

### OM permeability studies

OM sensitivity against SDS/EDTA was judged by colony-forming-unit (CFU) analyses on LB agar plates containing indicated concentrations of SDS/EDTA. Briefly, 5 mL cultures were grown (inoculated with overnight cultures at 1:100 dilution) in LB broth at 37 °C until OD_600_ reached ˜0.4-0.6. Cells were normalized by optical density, first diluted to OD_600_ = 0.1 (˜10^8^ cells), and then serially diluted (ten-fold) in LB broth using 96-well microtiter plates. 2.5 μL of the diluted cultures were manually spotted onto the plates, dried, and incubated overnight at 37 °C. Plate images were visualized by *G*:Box Chemi-XT4 (Genesys version 1.4.3.0, Syngene).

### SDS-PAGE, immunoblotting and staining

All samples subjected to SDS-PAGE were mixed 1:1 with 2X Laemmli buffer. Except for temperature titration experiments, the samples were subsequently either kept at room temperature or subjected to boiling at 100 °C for 10 min. Equal volumes of the samples were loaded onto the gels. As indicated in the figure legends, SDS-PAGE was performed using either 12% or 15% Tris.HCl gels (53) or 15% Tricine gel (54) at 200 V for 50 min. After SDS-PAGE, gels were visualized by either Coomassie Blue staining, or subjected to immunoblot analysis. Immunoblot analysis was performed by transferring protein bands from the gels onto polyvinylidene fluoride (PVDF) membranes (Immun-Blot 0.2 μm, Bio-Rad, CA, USA) using semi-dry electroblotting system (Trans-Blot Turbo Transfer System, BioRad). Membranes were blocked for 1 h at room temperature by 1 X casein blocking buffer (Sigma-Aldrich), washed and incubated with either primary antibodies (monoclonal α-MlaA (12) (1:3000) and α-OmpC (31) (1:1500)) or monoclonal α-His antibody (pentahistidine) conjugated to the horseradish peroxidase (HRP) (Qiagen, Hilden, Germany) at 1:5000 dilution for 1 – 3 h at room temperature. Secondary antibody ECL^™^ anti-mouse IgG-HRP was used at 1:5000 dilution. Luminata Forte Western HRP Substrate (Merck Millipore) was used to develop the membranes, and chemiluminescence signals were visualized by *G*:Box Chemi-XT4 (Genesys version 1.4.3.0, Syngene).

## Acknowledgements

We thank Swaine Chen and Varnica Khetrapal (Genome Institute of Singapore, A*STAR) for providing α-OmpC antibody, and reagents for negative selection and assistance in this technique. We also thank Michael Berne (Analytical Core Facility, Tufts Medical School) and Ross Tomaino (Taplin Mass Spectrometry Facility, Harvard Medical School) for performing Edman sequencing and MS/MS, respectively. We gratefully acknowledge computing resources provided by the National Supercomputing Center Singapore (http://www.nscc.sg). Finally, we thank Swaine Chen for critical discussions and comments on the manuscript. J.Y. was supported by the National University of Singapore Graduate School for Integrative Sciences and Engineering scholarship. J.K.M. and P.J.B. acknowledge support from the Singapore Ministry of Education Academic Research Fund Tier 3 grant (MOE2012-T3-1-008). All experimental work was supported by the National University of Singapore Start-up funding, the Singapore Ministry of Education Academic Research Fund Tier 1 and Tier 2 (MOE2013-T2-1-148) grants (to S.-S.C.).

## Author contributions

J.Y., K.W.T., D.A.H., P.J.B. and S.-S.C designed research; J.Y., K.W.T., Z.-S.C. performed all wet lab experiments described in this work; D.A.H. and J.K.M. performed all MD simulations; J.Y., K.W.T., D.A.H., P.J.B. and S.-S.C. analyzed and discussed data; J.Y., K.W.T., and S.-S.C. wrote the paper.

